# Metatranscriptomic Profiling Reveals Species-Level Microbial Shifts and Metabolic Remodeling in Feline Oral Inflammatory Disease

**DOI:** 10.64898/2026.03.08.710420

**Authors:** Claire A. Shaw, Maria Soltero-Rivera, Rodrigo Profeta, Cory L. Schlesener, Carol B. Huang, Angel Avalos, Boaz Arzi, Bart C. Weimer

## Abstract

Progressive oral mucosal inflammatory diseases are common among mammals. Cats are a valuable natural model for these conditions because they frequently develop oral diseases with varying severity, yet causative microbes remain unidentified in part because longitudinal studies are challenging and sampling is difficult. The lack of individual pathobionts suggests community-scale taxonomic and functional remodeling of the microbiome may be a contributor to oral disease. This study evaluated the microbiome composition and function of 33 cats across three cohorts with different levels of inflammation: healthy, aggressive periodontitis, and feline gingivostomatitis. Ultradeep metatranscriptomic sequencing was used to assign microbial taxonomy and examine functional changes via differential gene expression analysis to reveal genera that maintained stable relative abundance across all three disease states, including *Porphyromonas* and *Treponema*, while others had more subtle shifts in species activity, including multiple members of *Moraxella* and *Mycoplasmopsis.* Disease status was marked by co-occurring changes in low activity species accompanied by microbiome-level changes in protein, arginine, and nitrogen metabolism. Cats with aggressive periodontitis and gingivostomatitis displayed microbiome shifts in species that correlated to disease state. Microbial differential expression analysis revealed induction of stress-related genes and metabolic genes involved in amino acid metabolism, polyamine production, and nitric oxide production. Together, species identification and functional profiling suggest oral inflammation was correlated to shifts in the activity of multiple species that were involved with NO and polyamines. These coordinated metabolic signatures represent potential diagnostic targets for feline oral inflammatory disease.

## Background

Mammalian periodontitis, routinely seen in both humans and companion animals, is clinically important and is a significant impairment for animal health [1]. The etiology of periodontitis is complex, with previous work indicating oral inflammation stems in part from interaction between the host immune system and the oral microbiome [2]. Recently, oral microbes have been associated with systemic diseases that include neurodegeneration, colon cancer, and diabetes mellitus [3-5]. This increasingly recognized connection between oral and systemic health, combined with mounting evidence of microbial involvement, highlights a significant research gap. Surprisingly few studies have examined how oral inflammatory diseases relate to shifts in microbiome composition and function. Companion animals, such as dogs and cats, uniquely allow for sampling of a treatment-naïve population prior to the application of aggressive treatment strategies, as naturally occurring disease is routinely seen in clinics prior to intervention. The consistent clinical presentation of oral inflammatory diseases with varying severity makes cats an ideal population for investigating disease-related oral microbial shifts. Findings from these studies may then be applicable to other mammals.

Two important oral inflammatory diseases in cats are aggressive periodontitis (AP) and feline chronic gingivostomatitis (FCGS). Though progressive oral diseases, like AP and FCGS, are some of the most common culprits of oral tissue deterioration in cats, no single causative agent has been identified for either condition. Diagnosis is further complicated because the disease phenotypes can overlap somewhat as disease progresses [6], making the development of accurate diagnostic tools and effective treatments difficult [6]. The healthy oral microbiome of cats is dominated by species from the genera of *Porphyromonas, Treponema, Fusobacterium,* and *Moraxella* [7, 8]. Breed, age, diet, and indoor/outdoor status are known to affect the distribution of these genera in the microbiome [7, 9]. It is likewise important to note biogeographical niches, such as the tongue dorsum, buccal mucosa, and subgingival regions, also affects the microbial composition [8, 10]. These different oral niches are distinct in their microbiome composition and can hold different relevance to diverse oral diseases. However, profiling easily accessible samples such as buccal swabs, even when distal to the disease site, can provide clinically relevant information in a minimally invasive manner. [8]. Prioritized use of these minimally invasive sample types is critical for future widespread diagnostic use, as invasive samples are unlikely to be taken during routine clinical visits.

Though many factors alter the microbiome composition, oral inflammatory disease broadly has been identified as having a significant correlation with changes in the microbiome membership across the oral cavity [8, 11-15]. Previous work suggests that AP and FCGS communities are enriched with species from *Porphyromonas, Neisseria, Bacteroides, Treponema, Clostridium,* and *Fusobacterium* [13, 15, 16]. However, the use of low-resolution 16S rRNA sequencing used in some studies limits taxonomic interpretation to family or genus-level conclusions relative to disease. This has led to difficulty in assigning causality to the role of specific organisms to disease initiation and stages. These limitations suggest that more exacting methods looking at species or strain-level changes are necessary to devise accurate diagnostics and effective treatment strategies. While none of the previous studies on the diseases have identified a causal bacterial pathogen for either AP or FCGS, other work has suggested feline calicivirus, feline leukemia virus, and feline immunodeficiency virus, may be linked to the onset or exacerbation of FCGS [12, 15, 17]. Despite literature suggesting viral load contributes to FCGS manifestation, definitive causality between these viruses, the microbiome, and inflammatory status remains ambiguous.

When no clear pathobiont or viral agent emerges, as with feline AP or FCGS, researchers may look beyond taxonomic abundance or shifts in individual organisms. Partnering metabolic function within the community from specific organisms offers an additional informational dimension that can provide new insights for diagnostics and therapeutic options [18]. While studies evaluating functional (i.e. biochemical) activity of the oral microbiome are scarce in cats, studies in the human oral microbiome suggests a role for microbial nitrogen metabolism, specifically arginine catabolism, in preventing or exacerbating dental disease [19]. The arginine deaminase system in bacteria catabolizes arginine to produce ammonia and carbon dioxide, ultimately raising the oral pH and preventing demineralization of dental structures [20].

Bacterial community dynamics complicate the potentially protective role of arginine. These dynamics alter the metabolic landscape in ways that favor established dental pathogens, including *Porphyromonas gingivalis* [21, 22]. Co-cultures of *Streptococcus gordonii* and *Fusobacterium nucleatum* result in amino acid-rich environments that support the production of putrescine, a polyamine produced from arginine metabolism [22]. This metabolic crosstalk between oral bacteria leading to putrescine accumulation potentiates the growth of dental pathogen *P. gingivalis* and accelerates biofilm formation [22]. Bacterial production of polyamines also exerts direct consequences on host tissues through the modulation of inflammatory cascades [23]. Limited work details how arginine-derived polyamines from bacterial metabolism regulate oral inflammation. However, studies in other body sites, including the gut, suggest these bacterial metabolites regulate microbiome community membership dynamics and exert direct effects on local tissue that negatively impact health [23].

Arginine metabolism by bacteria in the oral cavity contributes to the formation of bioactive nitric oxide (NO) [24, 25]. Nitrate reduction by oral commensals contributes to systemic host health via production of NO, which interfaces directly with host cell signaling pathways that modulate blood pressure and inflammatory cascades [24-26]. Metabolic regulation of and by oral microbes may contribute to disease progression. However, the complex nature of metabolite crosstalk remains poorly understood and warrants further investigation into how bacterial metabolism at the epithelial interface contributes to oral inflammatory disease. Although arginine and its metabolites influence microbial community dynamics, the specific role of bacterial proteolytic metabolism in AP or FCGS onset remains unknown.

Understanding the role of oral microbial metabolism in chronic oral and dental diseases requires two key elements. First, researchers need knowledge of the taxonomic distribution and functional landscape of a healthy microbiome. Second, they need robust analysis methods to accurately identify species-level shifts that accompany these diseases. In this work, we use ultra-deep (>150 million reads/sample) total shotgun RNA sequencing (metatranscriptomics) to investigate the microbial community membership of three different cat populations; from healthy to aggressive periodontitis affecting the supporting structures of the tooth, to chronic gingivostomatitis affecting the keratinized and non-keratinized mucosa. Metatranscriptomics served a dual purpose in this study: 1) to identify microbiome membership, including viruses and down to the species/strain level for bacteria, and 2) determine the microbial activity including changes in bacterial metabolism at both the community and genus-level. This parallel analytic approach allowed us to test the hypothesis that species-specific microbial shifts and accompanying metabolic remodeling produce a distinct functional signature in the buccal mucosa of cats with AP and FCGS.

The disease-stratified cat populations revealed community rearrangement of bacterial species as disease progressed. Specifically, as we move from disease affecting masticatory mucosa and periodontal tissues to disease affecting not only that compartment but also lining mucosa, we observed a large reduction in *Moraxella* species with a co-occurring expansion of *Mycoplasmopsis* in FCGS-afflicted cats. Concurrent functional analysis uncovered the induction of bacterial genes involved in stress, arginine metabolism, and NO production were associated with progression from a healthy oral cavity to early onset disease and finally chronic and severe inflammation (Figure 1). These data suggest AP and FCGS are a disease stemming from coordinated reorganization of microbial composition and function that parallels tissue-specific disease.

**Figure 1.**
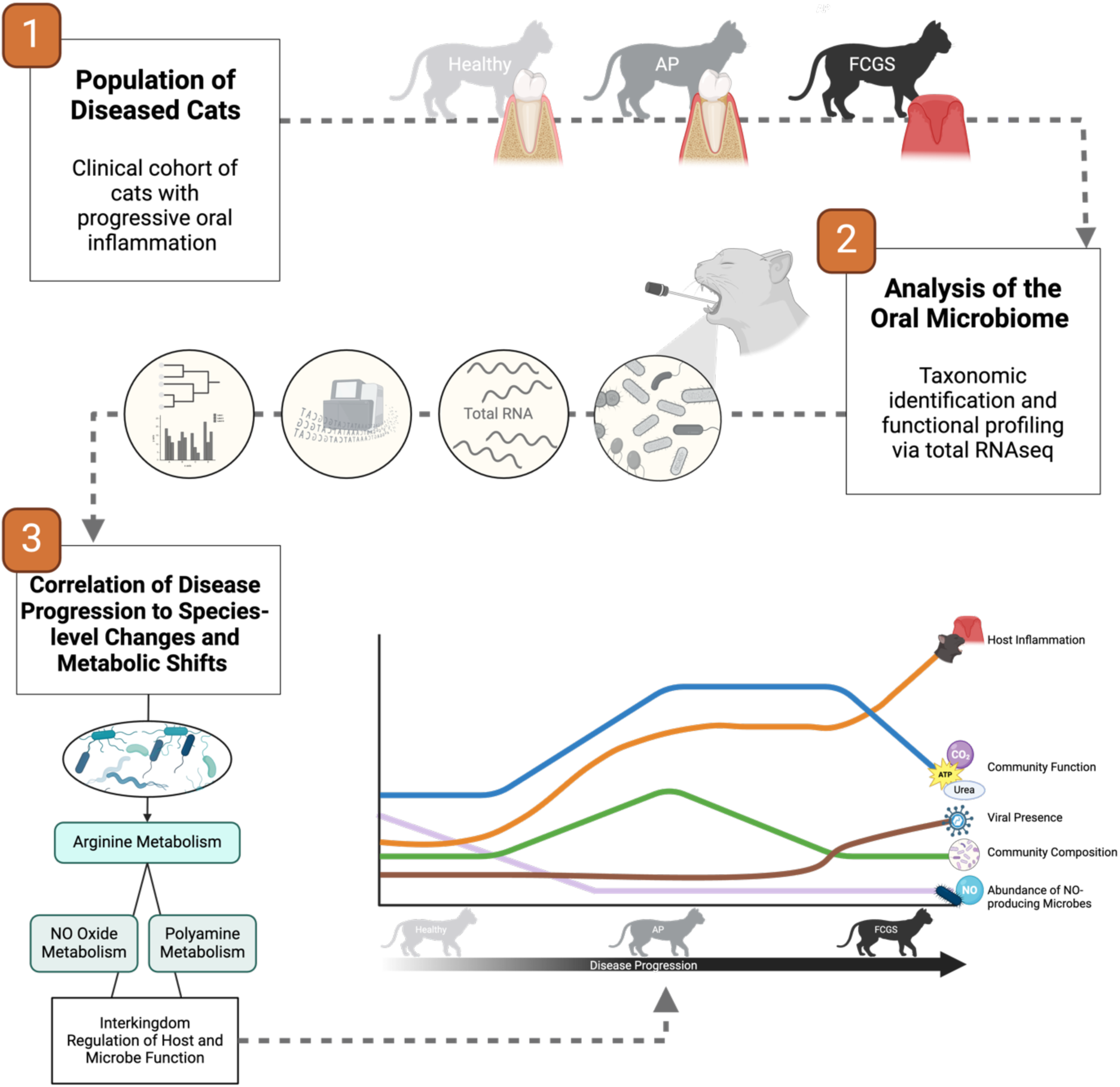
Graphical overview of the study design, sample processing, and biological observations

## Methods

### Patient Criteria and Sample Collection

A total of 33 cats were sampled for metatranscriptomics sequencing. Seven individuals had multiple samples collected for sequencing. Three cats had longitudinal samples obtained and for those the samples were categorized as per the disease present at the time of sample collection, and four cats had both left and right swabs collected. Supplemental Table 1 provides a breakdown of host age, sex and breed at the time of sampling. Privately owned cats presenting at the University of California Veterinary School were used in this study. All study procedures were reviewed and approved by the University of California-Davis Institutional Animal Care and Use Committee under IACUC #22738 and signed owner consent was obtained before sampling.

Cats were sampled and clinical definitions and criteria for diagnosis of aggressive periodontitis (AP) and feline chronic gingivostomatitis (FCGS), along with exclusion criteria, were based on clinical, radiographic, and histopathologic findings. For AP, cats ≤2 years old at initial presentation were included if they had moderate-to-severe inflammation of the gingiva that did not cross the mucogingival junction and supporting radiographic evidence of periodontitis [6]. For FCGS, diagnosis required moderate-to-severe inflammation lateral to the palatoglossal folds, gingivitis, radiographic evidence of periodontitis and resorption, as well as histologic confirmation [15]. Cats with oral neoplasia, osteomyelitis, or auto-immune diseases as well as cats with immunocompromising conditions such as diabetes mellitus or ongoing chemotherapy were excluded from the study.

All samples were collected by a Board Certified Veterinary Dentist™ (BA, MSR) during awake oral examination (18) or at the start of routine periodontal treatments (23) prior to rinsing or scaling the oral cavity. For each cat, a cytobrush (FLOQSwabs, Coplan, Italy, EU) was used to swab the caudal buccal mucosa. The swab was placed in a sterile conical tube that contained 500 μL of DNA/RNA Shield (Zymo, Irvine, CA, USA). The samples were vortexed before being frozen and stored at −20°C until processing. A visual overview of this study’s methodology can be found in Figure 2.

**Figure 2.**
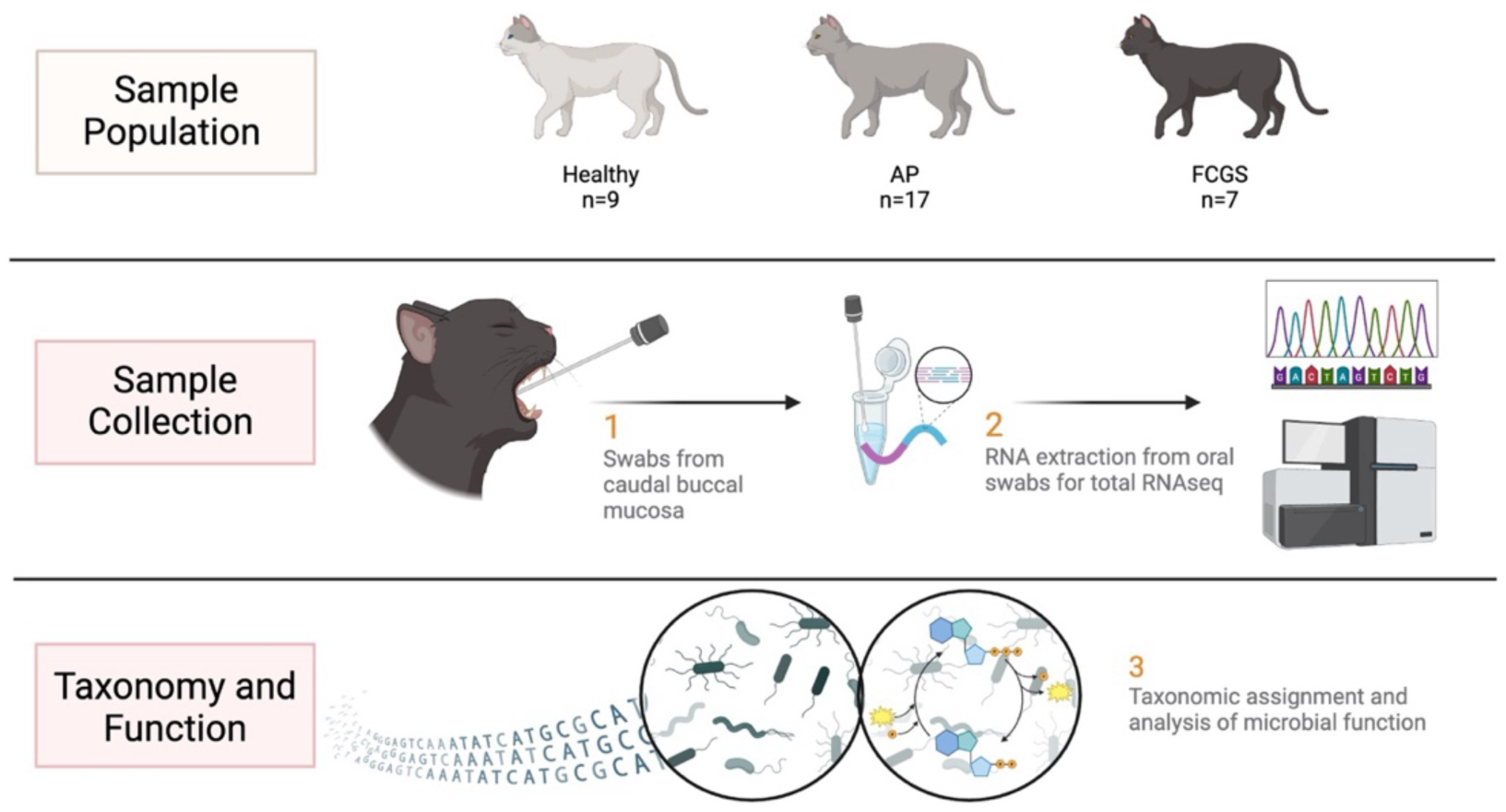
Method overview of the sampling population, sampling procedure and data analysis. The oral microbiome of 33 cats was sampled via swabbing the caudal buccal mucosa. Total RNA was extracted from each swab and sequenced. Sequencing data was then used to assign taxonomy at the species level and to develop functional profile for each oral microbiome.

### RNA Isolation and Library Preparation

Each sample was incubated in ice for 30 minutes after adding 1 mL of Trizol LS (Ambion by Life Technology, Carlsbad, CA, USA). After 30 minutes on ice, the swabs in Trizol were vortexed to transfer bacteria into solution and the swab was then removed from the tube. Bacterial cells were enzymatically lysed following the protocol used by the 100K Pathogen Genome Project for bacterial DNA isolation [27]. Briefly, samples were collected via centrifugation and suspended in Trizol LS. RNA was extracted from TRIzol LS (Ambion, Austin, TX, USA) following manufacturer instructions. RNA sequencing libraries were prepared as described previously with no rRNA depletion [28-30]. RNA purity (A260/230 and A260/280 ratios ≥ 1.8, ≤ 2.0) was done using the Nanodrop and integrity was confirmed with TapeStation (Agilent Technologies Inc., Santa Clara, CA, USA).

Sequencing libraries were constructed as previously described (REF) using the enzymatic based KAPA HyperPlus Library Preparation kit (KK8514) (Kappa Biosystems, Wilmington, MA, USA) on a PerkinElmer Sciclone G3 (PerkinElmer Inc. Waltham, MA, USA). Double-stranded cDNA were fragmented, indexed using Integrated DNA Technologies Weimer 384 TS-LT DNA Barcodes (Integrated DNA Technologies, Coralville, IA, USA), followed by dual-SPRI size selection and PCR amplification. Final library size was confirmed by PerkinElmer Labchip GX, HT D1K kit (PerkinElmer Inc. Waltham, MA, USA), with a targeted average fragment size of 300 to 450 bp. Library quantification was done using KAPA Library Quantification Kit (KK4824) (Kapa Biosystems, Wilmington, MA, USA) to ensure normalized concentration for sequencing pooling. Libraries were sequenced as 150 bp paired-end sequencing on an Illumina NovaSeq S4 (Illumina, San Diego, CA, USA).

### Taxonomic Assignment

RNA sequence analysis was done as described previously [30] using Trimmomatic (version 0.39) [31] to remove low-quality sequence and sequencing adapters, using settings “trimmomatic PE {input} {output} ILLUMINACLIP:{adapters}:2:40:15 LEADING:2 TRAILING:2 SLIDINGWINDOW:4:15 MINLEN:50”. Sequence data quality was reviewed with FastQC (version 0.11.9) [32]. Trimmed sequence reads were then separated between host and microbial content using Kraken2 (version 2.0.8) [33], using the “--classified-out” and “--unclassified-out” functions and standard settings (k-mer size = 35). For host separation a Kraken2 database was built with *Felis catus* reference genome RefSeq GCF 000181335.3 (version 9.0) [34].

Microbial sequence reads were assigned taxonomic identities with Kraken2 with microbial reference database, using standard settings (k-mer size = 35). The microbial database was a standard Kraken2 database build of NCBI RefSeq genomes, following the Kraken2 manual protocol (https://github.com/DerrickWood/kraken2/blob/master/docs/MANUAL.markdown, accessed on 13 May 2022), incorporating the categories archaea, bacteria, viral, fungi, protozoa, and UniVec Core (built/downloaded 13 May 2022).

Taxonomically-assigned reads were statistically proportioned to the respective taxa at the species level with Bracken (version 2.6.1) [35] according to previously used methods [28]. Braken species database was made from the Kraken2 database by standard protocol, using the parameters of k-mer size = 35 and read size = 150. Quality metrics for the sequencing data and taxonomic assignment can be found in Supplemental Table 2.

### Functional Annotation and Analysis

De novo transcriptome assembly was done using Trinity v2.15.1 [36], which reconstructs full-length transcripts from quality-trimmed FASTQ reads. Transcript abundance was quantified using Salmon v1.10.22 [37]. Differential expression analysis was conducted using DESeq2 [38], implemented within Trinity’s ‘run_DE_analysis.pl’ script, to compare cohort conditions (healthy vs. AP; healthy vs. FCGS; FCGS vs. AP). Resulting data were visualized in R (Version 4.2.3) using ggplot2 package to generate a volcano plot.

Functional annotation of assembled transcripts was performed using Prokka v1.14.6 [39] and Eggnog-mapper v2.1.12 [40]. Prokka, incorporating Prodigal [41], predicted coding sequences from the transcripts, while Eggnog-mapper provided orthology-based functional and evolutionary annotations using the EggNOG v5.0.2 database.

For genus-specific transcriptome profiling, quality-filtered and adapter-trimmed reads were taxonomically classified with Kraken2 (v2.1.3; standard database built on July 6, 2023). The ‘--classified-out’ option retained only reads assigned to known taxonomic identifiers. Reads corresponding to nine target genera (*Bacillus, Fusobacterium, Moraxella, Mycoplasmopsis, Neisseria, Porphyromonas, Streptococcus, Treponema,* and *Veillonella*) were extracted using a custom Bash script (grep -A 3 -w “kraken:taxid|${TAXID}” --no-group-separator) and compiled into paired FASTQ files for each genus. Each genus-specific set of reads was assembled and annotated following the same workflow described above.

Metabolic reconstruction and pathway completeness were assessed using anvi’o (v8) [42] through the anvi-run-kegg-kofams and anvi-estimate-metabolism modules, referencing the KEGG KOfam database. Pairwise ecological interactions between genera were modeled using RevEcoR [43], which calculates metabolic complementarity and competition indices based on predicted metabolic networks. Genus-by-genus scores were visualized as non-clustered heatmaps in ggplot2.

Pathway tools (Version 27.5) [44] in tandem with the MetaCyc Database [45] (SRI international, Menlo Park, CA, USA) were used to calculate enrichment scores and identify enriched GO groups at the microbiome community level. The presence of genes related to arginine metabolic processes in select oral microbes was evaluated using the “Species Comparison” function in MetaCyc along with publicly-available reference genomes for each organism. Gene expression data related to inducible nitric oxide synthase (iNOS) in the host was evaluated using datasets in healthy and FCGS-afflicted cats from a previous study by this paper’s authors [46] and visualized using Ingenuity Pathway Analysis (Qiagen, Redwood City, CA, USA).

### Analysis and Plotting of Microbiome Taxonomy

The relative abundance matrix from the Braken output was used to analyze the taxonomic distributions of each feline cohort; healthy, AP, and FCGS. The Shannon diversity index for each sample, then grouped into their respective cohorts, was calculated using the Vegan package in R (Version 4.2.3), assessed for significance with a two-way ANOVA test, and plotted using Prism (Version 10.2.2) (GraphPad, Boston, MA, USA) using a Bonferroni correction.

Species-level distributions across the healthy, AP, and FCGS cohorts were visualized using alluvial plots drawn in R (Version 4.2.3) using the ggplot2 and ggalluvial packages with aggregated abundance data for each cohort. The relative proportion of reads were used as the y-axis to represent relative abundance of genus and species. Strata on each alluvial plot were made to represent genera and alluvia represent species within each genus.

Correlation plots were constructed to visualize the relative abundances of various genera and species, based on data processed through Bracken analysis. Read counts were normalized by dividing the raw abundance of each organism by the total abundance within its corresponding sample, thus enabling the calculation of average abundances across three distinct treatment groups: healthy, AP, and FCGS. The dataset was further segmented for pairwise comparisons among the groups - Healthy vs. AP and Healthy vs. FCGS. Log_2_-transformed scatter plots were created using R (Version 4.2.3) with ggplot2. Final versions of the plots were labeled manually in Inkscape (Version 1.0) accessed via GitHub (https://github.com/inkscape/inkscape).

Correlation networks of microbial genera and species within each cohort (healthy, AP, and FCGS) were assessed using Spearman’s correlation coefficient calculated using the abundance matrix from Braken in R (Version 4.2.3) with the Hmisc package. Cloud networks of genera correlations were visualized using Cytoscape (Version 3.10.2) and species-level heatmaps were visualized in R (Version 4.2.3) using the corrplot package.

## Results

### Species Switching is Associated with Aggressive Periodontitis and Gingivostomatitis

To define the community structure in each disease status we examined the taxonomic diversity of each cohort, between healthy, AP, and FCGS (Figure 3). While all cohorts surveyed shared 6,193 species and strains, the two inflammatory disease cohorts (AP and FCGS) displayed increased numbers of unique microorganisms compared to healthy subjects (Figure 3A). The total diversity and number of shared and unique species in the cohorts were likewise reflected in non-significant (p>0.05) differences in the Shannon Diversity metric across all groupings (Figure 3B).

**Figure 3.**
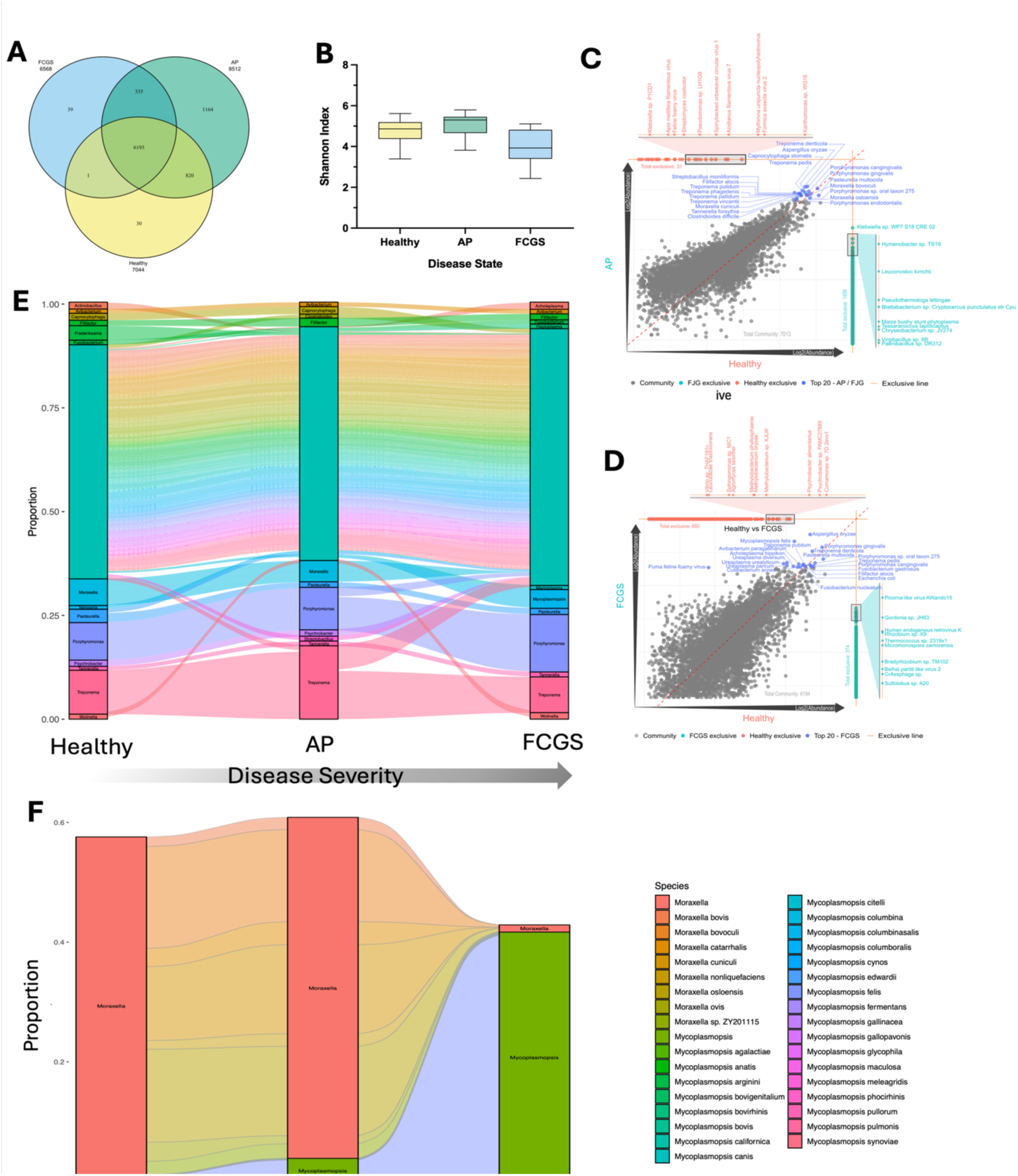
Changes in the taxonomy of the oral microbiome across healthy and disease at the genus and species level. Comparisons were made between a healthy feline cohort, those with unrelated oral disease (TR), early inflammation from aggressive periodontitis (AP), and severe inflammation from feline gingivostomatitis (FCGS). (A) Venn diagram of the unique and overlapping species between all three cohorts. (B) Alpha diversity of the feline cohorts as measured by the Shannon Index. (C & D) Correlation plots displaying the shared and unique species between AP and healthy felines (C) or between FCGS and healthy felines (D). Each axis represents a cohort and each dot represents a single species, while movement along each x-y-axis illustrates increasing Log2 abundance of species in the respective cohort. Dots on the center line are in equal abundance in both cohorts. Dots found on the lines in the far right or upper edges of the plot are unique to one cohort in the comparison. Species of interest have been individually labeled. (E) Alluvial plot illustrating species switching from the healthy microbe through early inflammation (AP) and to severe inflammation (FCGS). Strata are comprised of genera with the y-axis representing the proportion reads from each genera, indicating relative abundance. Genera with a total proportion of reads less than 0.1 were collapsed into the teal stratum. Alluvia represent species in each genera. (F) Subset alluvial plot illustrating the change in abundance of *Mycoplasmopsis* and *Moraxella* species across health and disease.

Considering the number of shared species and relatively minimal difference in diversity across cohorts, species-level differences in the taxonomic distributions by disease groups were investigated (Figures 3C & 3D). Relative abundance (i.e. activity) values for each species in the combined cohort microbial profile were plotted and revealed many common oral species were shared between diseased and healthy cats at similar relative abundance (e.g. *T. denticola* and *P. gingivalis*) (Figures 3C & 3D). More broadly, the close clustering pattern of dots denoting organisms around the center line in both the AP-Healthy and FCGS-Healthy comparison suggests a high level of similarity in the oral microbiome composition regardless of disease status, supporting the lack of significant changes in the Shannon Diversity Index (Figure 3B).

Investigation of taxonomic changes across these inflammatory diseases revealed relative abundance was altered at the species-level (Figure 3E). *Avibacterium, Filifactor, Porphyromonas, Tannerella,* and *Treponema* were present at similar abundance within each of the three cohorts. Other genera, such as *Actinobacillus* and *Neisseria*, were greatly reduced (total proportion of reads <0.1) in the AP group compared to health. In contrast, *Capnocytophaga* and *Psychrobacter* both remained above the total proportion of reads <0.1 in Healthy and AP but collapsed to relative abundances below this mark in FCGS. In FCGS, low abundance genera in AP including *Acholeplasma, Mycoplasmopsis,* and *Marinplasma* expanded to above the low 0.1 proportion cutoff and others, like *Moraxella,* greatly reduced in abundance.

Seemingly stable genera like *Porphyromonas* and *Treponema*, experienced within-genus species shifting (Figure 3E, Supplemental Figure 1). In addition to bacterial shifts, the presence of four select feline viruses in healthy, AP and FCGS affected cats was examined (Supplemental Figure 2). The abundance of feline calicivirus and feline foamy virus increased modestly between AP to FCGS cats, while feline leukemia virus went from present in healthy and AP to absent in FCGS. The most notable change in viral abundance was with feline immunodeficiency virus, which nearly doubled in abundance in FCGS as compared to Healthy or AP cohorts.

Notably, *Moraxella* and *Mycoplasmopsis* were two genera that, while present in relatively low abundance, clearly showed altered proportions between healthy, AP, and FCGS subjects (Figure 3F). While *Moraxella* retained an equal proportion of the healthy and AP microbiome, with stable species distribution across the two groups, the genera were drastically reduced in the FCGS subjects. *M. bovoculli*, *M. cuniculi,* and *M. osloensis* were all uniformly proportioned in the healthy and FCGS samples, while *M. osloensis* became the dominant species of *Moraxella* in FCGS. In contrast to the noted decrease of *Moraxella* in FCGS, *Mycoplasmopsis* increased 4-fold from the healthy and AP groups to the FCGS group. *M. felis* was the dominant species of *Mycoplasmopsis* in healthy and AP and remained the dominant species in FCGS albeit with increased abundance.

### Bacterial and Viral Co-occurrence is Specific to Disease Status at Genus and Species Level

The observation that species-switching through disease state stratifies and potentially contributes to the extent of inflammation and suggests community dynamics within the microbiome are likely contributors to taxonomic distribution. To address this, the abundant genera (proportion > 0.1; Figure 3E) were examined for co-occurrence with the multiple taxonomies and with viral presence. In healthy cats, the abundance of feline calicivirus was not significantly correlated with any major genera in the oral microbiome of healthy cats (Figure 4A). In contrast, the presence of feline calicivirus was correlated with the most abundant genera in AP and FCGS cohorts (Figure 4B & 4C). Intriguingly the major genera were almost all positively correlated with each other, indicating strong community interaction and genus-genus interactions that shift in concert with AP and FCGS presentation.

**Figure 4.**
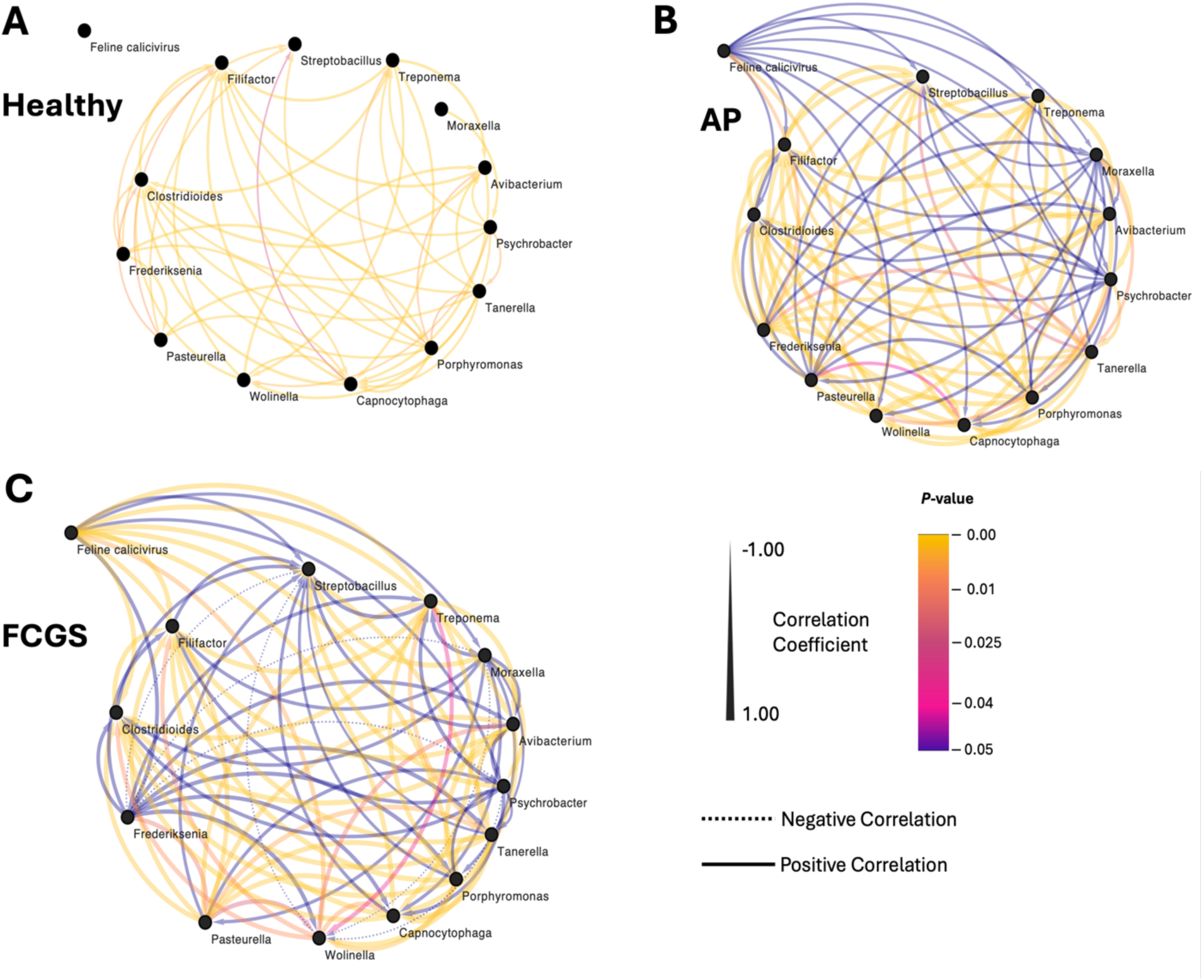
Co-occurrence networks of prominent genera in healthy (A), aggressive periodontitis (AP) (B), and feline chronic gingivostomatitis (FCGS) (C) oral microbiomes. Genera in this network analysis were picked from the stratum in Figure 2E and represent the most abundant genera in the three cohorts. Feline calicivirus was additionally included in this analysis as it has been suggested to be associated with FCGS. Spearman’s rank correlation coefficient was used to determine correlation values between genera based on relative abundance of each genera. Solid lines represent a positive correlation, dotted lines represent a negative correlation, and the absence of a line indicates no or nonsignificant correlation between genera.

Cementing that species-switching and the concurrent effects on the microbial community are more strongly associated with disease status than genera-level observations, co-occurrence of feline calicivirus and *Mycoplasmopsis* is highly species-specific (Supplemental Figure 3). Some *Mycoplasmopsis*, including *Mycoplasmopsis synoviae* and *M. felis*, show weak or no correlation to feline calicivirus load while others like *Mycoplasmopsis pulmonis* were significantly correlated with total viral load. This species-specific correlation to viral occurrence was likewise reflected among other bacterial species seen in this study.

### Metabolic Functional Shifts were Unique to Disease Stage

The species-specific pattern of co-occurrence suggests that functional microbial dynamics within the community may explain the role of the microbiome response linked to disease distinction. To assess functional shifts associated with each disease state, gene expression at the community level was compared between either AP or FCGS and Healthy. This analysis of differentially expressed genes within the total microbiome indicates that aggressive periodontitis (AP) and gingivostomatitis (FCGS) share some basic functions compared to healthy microbiome, but also notably differ in others like those related to stress responses (Figure 5).

**Figure 5.**
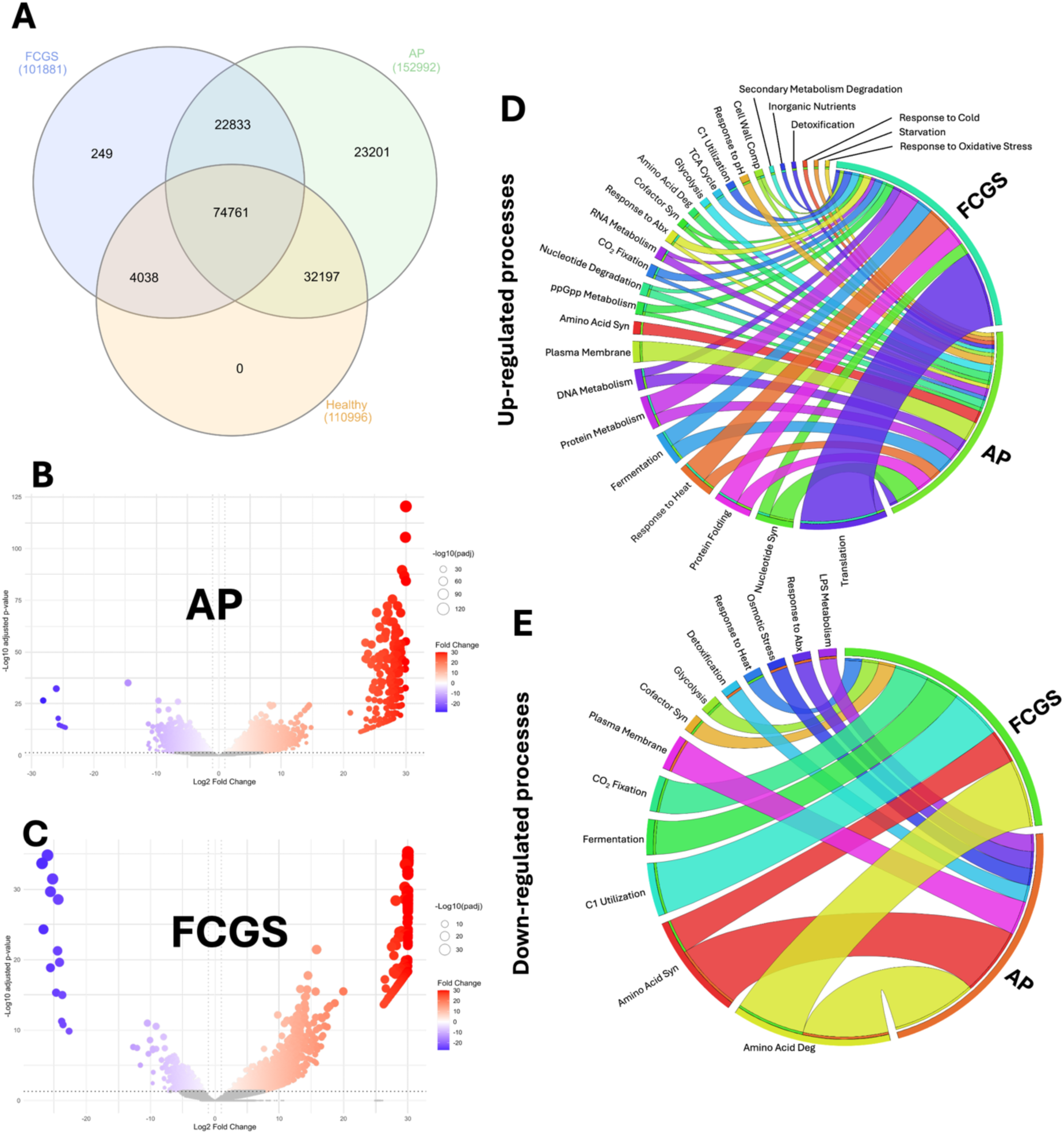
Functional profiles at the community level of the oral microbiome in aggressive periodontitis (AP) and feline chronic gingivostomatitis (FCGS). (A) Venn diagram representing the shared and uniquely expressed genes across healthy, AP, and FCGS oral microbiomes. (B & C) Volcano plots illustrating the significantly (adj-P val ≤ 0.05) up-regulated genes in red and significantly down-regulated genes in blue of the AP and FCGS microbiomes as compared to gene expression in the healthy microbiome cohort. (D & E) Chord diagram displaying the significantly (adj-P val ≤ 0.05) up- and down-regulated functions at the community level in AP and FCGS as compared to the healthy cohort. Line thickness corresponds to the enrichment (D) or depletion (E) score for each function, with thicker lines indicating a higher score.

While healthy cats had no uniquely expressed genes compared to disease states, microbes in the AP cohort differentially expressed 23,201 microbial genes not found in either the healthy or FCGS metatranscriptome (Figure 5A). In contrast the FCGS microbiome expressed 249 unique genes and showed the greatest overlap with AP expression at 22,833 shared genes. In addition to having the most uniquely expressed genes, the microbiome of AP animals primarily showed induction of gene expression, as noted by the primarily up-regulated significant genes, as compared to healthy animals (Figure 5B), and in comparison, to the mixed expression profile in FCGS (Figure 5C).

Microbes in the AP cohort communally up-regulated stress responses related to starvation, oxidative stress, cold, pH, and antibiotic pressure in comparison to healthy microbiomes, while FCGS microbial communities only displayed a response to antibiotic pressure (Figure 5D). AP communities also repressed some of these stress responses, like response to antibiotic pressure (Figure 5E), indicating that different members of the microbial community were differentially responding to environmental pressures. Though the response to specific stressors at the community level differs between AP and FCGS, both microbial communities induced ppGpp metabolism, a master regulatory molecule that controls the stress response in bacteria by regulating metabolism, persistence, and resistance in response to multiple environmental stressors [47].

In addition to stress responses, metabolism is another broad function that was differentially regulated across AP and FCGS microbial communities. Protein metabolism was significantly induced and enriched within the microbiome in both AP and FCGS cohorts (Figure 5D). Amino acid degradation was notably up-regulated only in the AP condition, while amino acid synthesis was both up- and down regulated in AP and only down-regulated in FCGS (Figure 5D & 5E).

### Protein Metabolism by Host and Microbiome Associated with Inflammation and Disease

The collective induction of biotic stress conditions combined with the noted induction of microbial amino acid biosynthesis in only AP led us to hypothesize that changes in microbial metabolism are associated with changes in the host inflammatory response. Arginine has previously been shown in human dental literature to be a bioactive amino acid that prevents tooth demineralization and regulates microbial community dynamics [48, 49]. Further enrichment analysis of genes positively expressed in the AP and FCGS cohorts, compared to healthy, revealed enrichment for the metabolically linked pathways of arginine, nitrogen, and polyamine metabolism at the community level (Figure 6).

**Figure 6.**
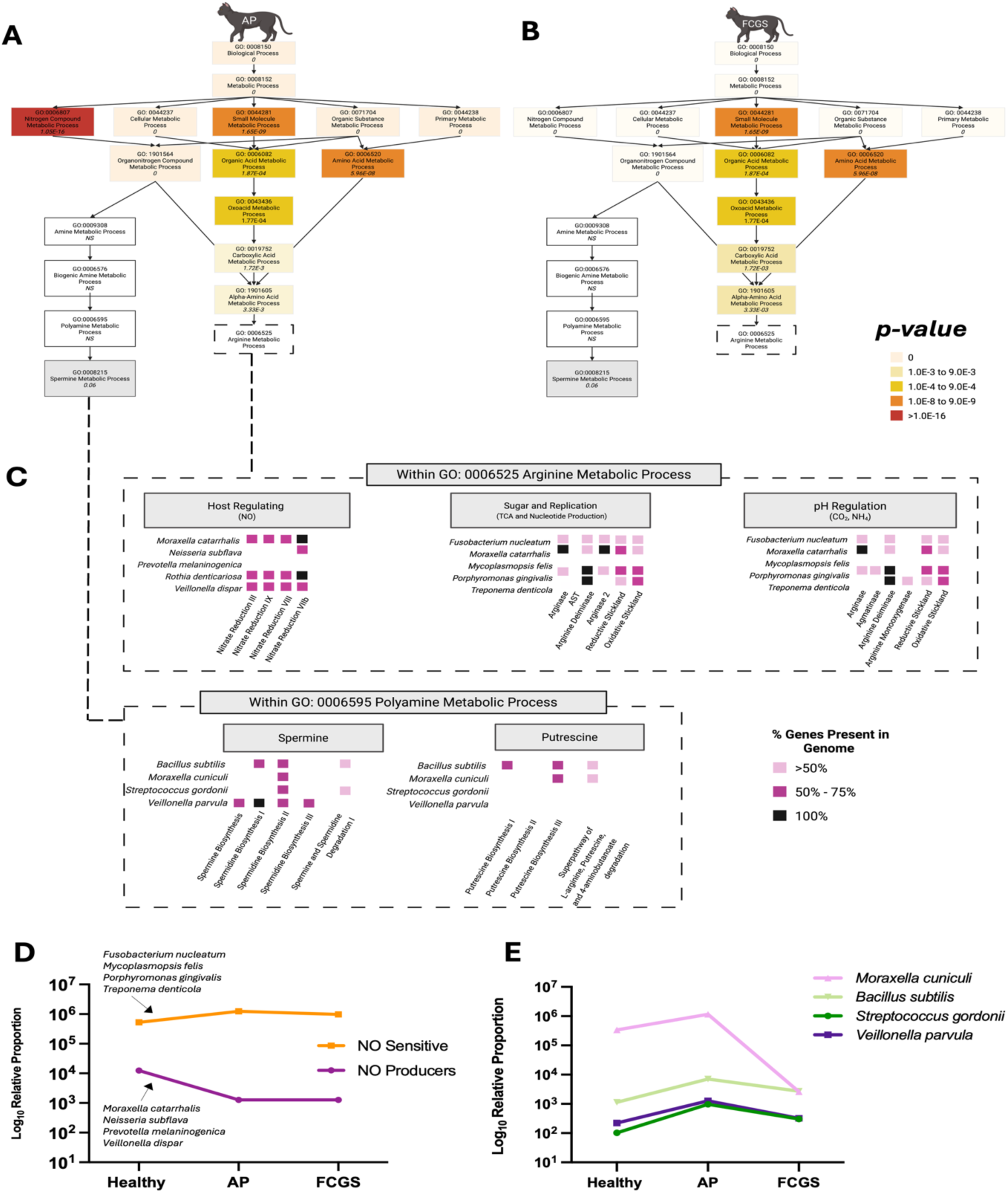
Arginine, nitrogen, and polyamine metabolic regulation across AP and FCGS cohorts. Enriched GO groups related to arginine metabolism in AP (A) and FCGS (B) cohorts as compared to the healthy cohort. All displayed GO identifiers are enriched in both disease groups compared to healthy. Color gradation corresponds to the *p-value* derived from Fisher’s test for significance with darker orange corresponding to more significance. (C) The presence of genes related to arginine or polyamine metabolism in selected oral bacteria were evaluated using reference genomes available in MetaCyc and broken down by metabolic pathway. Each square denotes presence of some or all genes in each pathway with color indicating percentage of genes present in that organism. Darker colors indicate more pathway coverage by gene presence in the genome. (D) Average relative abundance of the select microbes from panel C split by those able to produce nitric oxide (NO) and those sensitive to and unable to produce NO. (E) Shifting average relative abundance of microbial species relevant to polyamine metabolism from panel C.

The enrichment of gene ontology (GO) groups related to arginine metabolic processes (GO:0006525) was evaluated in AP and FCGS cohort and contrasted to healthy cats (Figure 6A & Figure 6B). Nitrogen compound metabolic processes (GO:0006807) was enriched in FCGS cats (*p-val* = 0) but was significantly enriched in AP cats (*p-val* = 1.05E-16) (Figure 6A & Figure 6B). Other bioactive arginine-derivative, polyamines, also had enriched, though non-significant, metabolic pathways in both disease states as compared to healthy (Figure 6A & Figure 6B).

Common oral bacteria with characterized arginine metabolism that displayed only minor abundance shifts in these cats include suspected cancer correlate *F. nucleatum* [3, 50], periodontal pathogen *P. gingivalis* [21, 51], and the highly proteolytic bacterium *T. denticola* [52]. The genetic capacity for arginine metabolism in these organisms was evaluated, along with two species that contrastingly displayed relatively large shifts in abundance, *M. catarrhalis* and *M. felis*. For polyamine metabolism, the genomic capabilities of *Bacillus subtilis, M. cuniculi, S. gordonii,* and *Veillonella parvula* were examined (Figure 6C).

The NO-producing bacteria carried genes for most nitrate reduction pathways, suggesting strong potential for nitrate reduction and NO production across the microbiome. In contrast, analysis of arginine pathways, which are linked to bacterial growth through nucleotide and sugar synthesis or to pH-regulating metabolites such as NH_₄_^⁺^ and CO_₂_, revealed more variable metabolic potential among individual species (Figure 6C). Few oral species possessed complete arginine metabolism pathways, and most showed incomplete polyamine pathways; however, all contained genes for spermidine production (Figure 6C). Fully understanding disease-associated shifts likely requires a holistic view that integrates host immune and metabolic interactions.

Nitrogen metabolism is complex and tightly regulated in both bacteria and animals. To assess whether bacterial–viral interactions may contribute to disease, we focused on nitric oxide (NO) production from arginine, a metabolite important to oral health [53-56] and produced by both bacteria and the host. We examined the metabolic capacity (Figure 6C) and abundance (Figure 6D) of nitrate-reducing bacteria including *Neisseria subflava*, *Prevotella melaninogenica*, *Rothia dentocariosa*, and *Veillonella dispar* [57]. These NO-producing bacteria showed an approximately 10-fold decrease in abundance collectively from healthy to AP conditions and remained low in the FCGS cohort. In contrast, NO-sensitive bacteria increased modestly across disease states (Figure 6D).

Polyamine-producing bacteria showed the opposite pattern: their relative activity increased from healthy to AP but slightly declined from AP to FCGS (Figure 6E). The limited change in NO-related bacteria between AP and FCGS suggests greater host influence on the microbiome as inflammation and immune activation become more widespread. Consistent with this, the NO-related NFκB inflammatory pathway was strongly induced in FCGS cats compared with healthy controls (Supplemental Figure 4A). Activation of this pathway, including the NO-producing NOS2–Calmodulin complex, indicates host-derived NO may have suppressed NO-sensitive microbes (Figure 6D) as bacterial NO producers declined.

Host polyamine metabolism displayed a more variable pattern (Supplemental Figure 4B). Predicted concentrations of ornithine, putrescine, and spermidine decreased in FCGS cats based on host gene expression, whereas spermine levels were predicted to increase. Correspondingly, the host spermine transporter SLC3A2 was upregulated, consistent with enriched bacterial spermine metabolism. These findings suggest spermine may act as an interkingdom metabolite influencing both host and microbial activity.

To further characterize microbial interactions, we used KEGG-based metabolic modeling to evaluate competition and complementarity among bacterial pathways (Figure 7A and 7B). Overall, competition scores were higher and more broadly distributed than complementarity scores, indicating overlapping metabolic capabilities across taxa. *Mycoplasmopsis* and *Neisseria* exhibited strong competitive interactions with multiple genera, whereas *Bacillus* showed minimal overlap or complementarity, suggesting more distinct metabolic niches. These results indicate that metabolic competition is a dominant feature of the oral microbiome, with selective complementary interactions that may help maintain community structure and contribute to disease-associated shifts.

**Figure 7.**
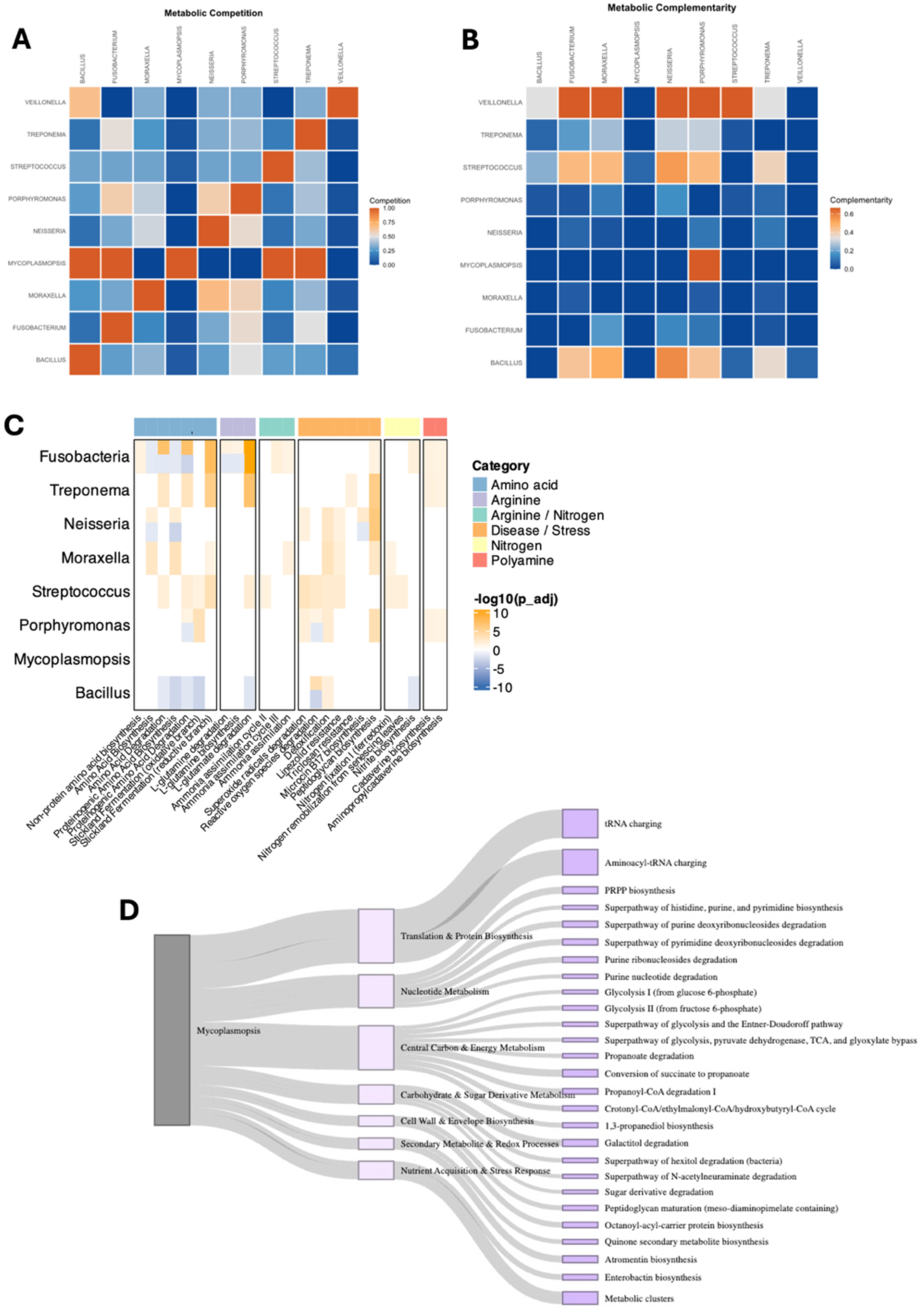
Metabolic competition and cooperativity in oral microbial genera observed in healthy and aggressive periodontitis (AP) microbiomes. Predicted microbial competition (A) and complementation (B) from KEGG-based metabolic networks for microbial genera from AP microbiomes. A higher metabolic competition score, illustrated by dark orange, suggests the genera have similar auxotrophic requirements, while metabolic complementation suggests metabolite sharing between genera to support auxotrophic requirements. (C) Gene set enrichment analysis of arginine, nitrogen and polyamine metabolic pathways in AP genera compared to healthy. Darker orange rectangles represent more significantly enriched pathways, while darker blue indicate repression. This analysis encompasses individual species variation, so multiple genera have mixed expression of pathways, as indicated by shared orange/blue rectangles. (D) Gene set enrichment of *Mycoplasmopsis* genes showing increased differential expression in AP compared to healthy microbiomes, with thicker lines corresponding to increased enrichment scores. In both (C) and (D) MetaCyc pathways were used as the background for enrichment with Bonferroni Correction and a p-value cutoff of 0.05.

To further investigate these relationships, we performed gene set enrichment analysis focusing on arginine, nitrogen, polyamine, and disease-associated pathways to assess the induction and repression of metabolic functions in the AP microbiome compared with healthy controls. This analysis revealed distinct metabolic trends in aggressive periodontitis (AP) compared to healthy cats (Figure 7C). Several genera, including *Fusobacteria*, *Treponema*, and *Neisseria*, showed enrichment of arginine and nitrogen-associated pathways, consistent with increased potential for nitric oxide and amino acid metabolism in disease. In contrast, Bacillus exhibited significant depletion of these pathways, suggesting reduced contributions to nitrogen or polyamine metabolism, or perhaps indicating use of exogenous metabolites. Pathways related to stress response and host interaction, including arginine–nitrogen overlap categories, were broadly induced across multiple genera, highlighting a community-wide metabolic adjustment either contributing to or responding to inflammation. These results indicate that shifts in nitrogen and arginine utilization distinguish AP-associated microbiomes from healthy controls and may reflect divergent ecological strategies among oral taxa.

Although *Mycoplasmopsis* did not show significant enrichment for these nitrogen-related pathways, it was enriched for a diverse set of central metabolic and biosynthetic processes, including tRNA charging, nucleotide biosynthesis and degradation (purine and pyrimidine superpathways), glycolysis, and short-chain fatty acid metabolism such as propanoate and succinate conversion (Figure 7D). Additional enrichment in sugar and hexitol degradation, peptidoglycan maturation, and secondary metabolite biosynthesis pathways (e.g., quinone, atromentin, and enterobactin) suggests that *Mycoplasmopsis* may sustain growth and competitive fitness in the inflamed oral environment through metabolic flexibility.

Together, these observations and collective host-microbe expression data suggest that arginine-related metabolic processes, specifically those leading to nitric oxide and polyamines, guide interkingdom interaction between host and oral microbes (Figure 8). The enriched arginine, nitric oxide, and polyamine metabolic pathways in microbiomes of diseased cats contributes to the circulation of bioactive compounds, which then exert effects on interconnected pathways related to host inflammation and health, like NFkB. The differential regulation of these host pathways and environmental pressures created can then act to regulate microbial abundance, creating a cycle of microbiome-host regulation that contributes to oral inflammation.

**Figure 8.**
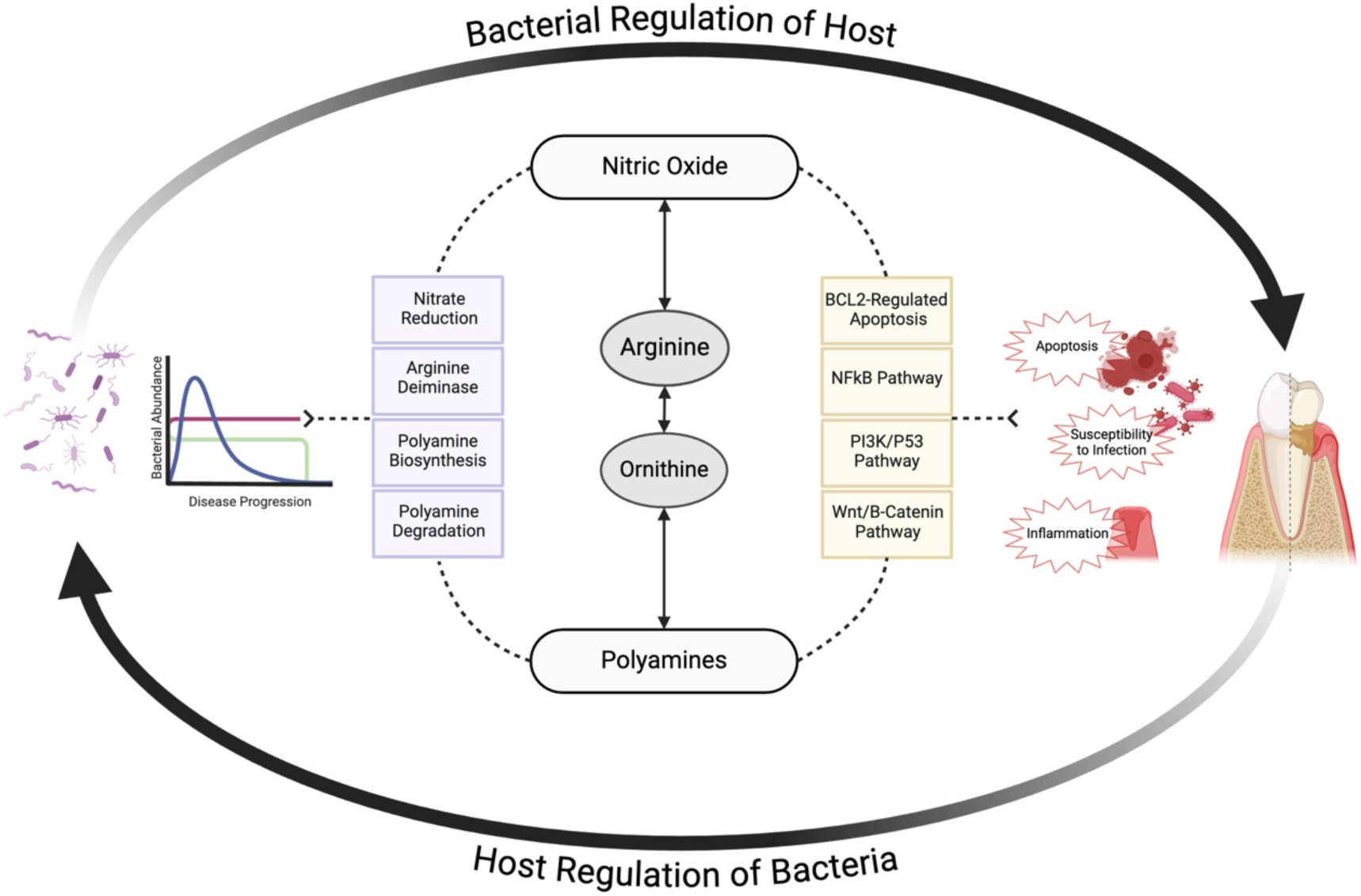
Interconnected arginine metabolism by both host and the oral microbiome regulate host expression of inflammation-related pathways and modulate microbial abundance. The bacterial metabolism of arginine to bioactive nitric oxide and polyamines affects host expression of related metabolic and inflammatory pathways. In turn the regulation of these inflammatory, apoptotic, and immunosuppressing canonical pathways exerts control over microbial activity and abundance in the oral cavity. This interkingdom interaction with arginine at the center creates a cyclic interaction in which microbial abundance and activity *affect* host expression which in turn modulates oral microbial presence and function. Bacterial pathways related to arginine metabolism and involved in polyamine or nitric oxide production are in purple while host pathways regulated by or involved in arginine metabolism are in yellow.

## Discussion

There is increasing clinical and scientific interest in the role oral microbes play in oral inflammatory diseases, but to date no study has chronicled the microbial landscape across oral diseases in cat cohorts using annotation accurate to the species level. In this study we sought to remedy this through the application of ultra-deep total shotgun metatranscriptomics in a population encompassing healthy, aggressive periodontitis and gingivostomatitis cats. The application of high-resolution taxonomic annotations to this feline population confirmed previous work showing genera-level abundances remain relatively steady across healthy and disease-related microbiomes [13], and confirming that use of low resolution techniques would miss these significant changes associated with disease.

AP and FCGS remain understudied, with limited treatment options available. Investigating the microbial contributions to these diseases is therefore imperative for improving clinical outcomes [6, 15]. The utilization of shotgun metatranscriptomics in tandem with species and strain identification to name functionally active microbes in the oral cavity of cats revealed previously unseen shifts at the species level in the oral microbiome. Previous studies identified *Moraxella* and *Mycoplasmopsis* as genera that shift in abundance during oral disease. Our study confirms these genus-level changes but reveals novel species-level variations within these genera that correlate with inflammatory disease state. Literature investigating the oral microbiome of cats across both healthy and diseased populations have primarily used 16S rRNA to name bacterial inhabitants of the oral cavity, and other amplicon-based approaches to identify fungi and viruses [8, 11, 58, 59]. Such studies have provided an observational understanding of the broad taxonomic distribution of the feline oral cavity across different contexts and identified actinobacteria, bacteroides, proteobacteria, firmicutes, and fusobacteria as some of the most consistently prevalent phyla in oral microbiomes [8, 58]. The application of ultra-deep metatranscript sequencing enabled microbial identification down to the species and strain level [60]. This approach also identified metabolically active microbes in the microbiome, thereby providing a more functionally relevant picture of the microbial community than previous studies using DNA-based methods. This study is, to the best of our knowledge, the first to employ metatranscriptomic profiling of the functional oral microbiome with progressing oral inflammation. This approach enabled high resolution observations of community dynamics coupled to host changes in oral inflammatory disease.

Common genera identified across the healthy and diseased cohorts in this study were *Porphyromonas*, *Treponema, Pasteurella,* and *Tannerella*, with gram-negative anaerobes of *Porphyromonas* being among the most stably abundant microbial populations. *P. gingivalis* more specifically has gained notoriety in the literature as a potential pathogen of the mammalian oral cavity [61, 62] and studies demonstrate that this opportunistic pathogen encodes virulence factors that induce local inflammation in the host [63]. The ability of *P. gingivalis* to disrupt the local environment has led to the suggestion that the macroscopic symptoms associated with *P*.

*gingivalis* are in part due to the microbiome remodeling that accompanies environmental shifts that alter metabolic pressures in microbes forcing them to respond with general stress-mediated pathways [62]. Intriguingly, this study found *P. gingivalis* was not only present in all populations, including healthy animals, but was similar in abundance across cohorts. While this finding suggests that outgrowth of *P. gingivalis* is not the causative agent in AP or FCGS, it doesn’t rule out that *P. gingivalis* activity could still be driving metabolically induced and subtle shifts in the microbial population.

One of these subtle, but notably impactful alterations of microbial composition across AP and FCGS cohorts, was the changes in *Moraxella* and *Mycoplasmopsis* populations. Though the relative abundance of these groupings in absolute value was comparatively low, *Moraxella* abundance collapsed in the FCGS cohort compared to healthy or AP. *Moraxella* are known inhabitants of the mammalian upper respiratory tract and were long thought to be commensal residents, but ongoing research suggests *M. catarrhalis* may be an opportunistic pathogen that exacerbate chronic conditions such as chronic obstructive pulmonary disease [64, 65]. Given the anatomical structure of the oral cavity and upper airways, it is unsurprising to observe overlapping respiratory and oral microbes [66]. Complicating the role of *Moraxella* is research suggesting high *Moraxella* carriage in the nasal cavity of human children is correlated with reduced COVID-19 infection [67], suggesting a potential protective role for *Moraxella* that was attributed to the bacteria’s regulation of amino acid metabolism in the microenvironment [67]. This mechanistic theory requires further validation. Nevertheless, it suggests how the collapse in *Moraxella* abundance and corresponding increase in *Mycoplasmopsis* may mark the severe inflammation characteristic of the FCGS population [18, 68].

Differential regulation of microbial amino acid metabolism may contribute to the deterioration of tissues seen in AP and FCGS [53, 69]. Though metabolites were not directly investigated during this study, functional annotation of the microbial community allowed insight into how microbial metabolism was differentially regulated across disease state. *M. catarrhalis*, along with other *Moraxella* species, can reduce available nitrogen in the oral cavity through assimilation via glutamate dehydrogenases activity that participates in concert with nitrate and nitric oxide reductases to modulate the production of NO [70]. Nitric oxide is a potent nitrogenous compound with concentration-dependent systemic effects in the host that include modulation of inflammatory pathways [26]. Multiple studies have previously associated dysregulation of NO production in the oral cavity with periodontitis and systemic diseases of the cardiovascular system and intestinal tract [71, 72]. The drastic reduction of *Moraxella* in the FCGS population suggests increased availability of nitrogen for use by neighboring microorganisms that have a robust proteolytic system, like *Mycoplasmopsis*. *Mycoplasmopsis* remains an understudied prokaryotic group in the oral cavity, but research suggests that this opportunistic airway pathogen shows the ability to induce NO dysregulation in the host through interference with immune-related signaling cascades [73, 74]. Aggressive periodontitis and FCGS are immune-mediated diseases marked by severe inflammation like that seen induced by nitric oxide modulated pathways [15, 75]. One potential microbially driven mechanism for inflammation in the oral cavity lies in the regulation of nitrogen and polyamine metabolism by the microbiome, leading to the production of inflammation-inducing levels of NO and increased circulation of bioactive polyamine spermine. As AP and FCGS are of unknown etiology and given the nuanced microbial shifts observed here, it remains that complex shifts and accompanying functional changes in the microbiome are in part responsible for observed symptoms.

The oral microbiome is thought to be a key player in maintaining oral health across species [10, 76, 77]. However, the specific role these microorganisms play in the onset or exacerbation of disease in mammals remains unclear. Notably, AP and FCGS mirror the inflammatory landscape of similarly debilitating oral conditions in humans [78, 79], presenting a potential unique opportunity to study human oral diseases in an accessible and clinically-relevant animal model, generating findings relevant to veterinary and human medicine alike. The cat genome shares >90% similarity with that of humans [80], indicating potential functional overlap in host genome-microbiome interactions. AP and FCGS are already present in the cat population and routinely seen in veterinary practice, thus reducing the need for invasive disease induction measures required by other animal models [81, 82]. Additionally, the complexity of FCGS increases the difficulty of developing a laboratory model with a high degree of fidelity, further supporting the importance of sampling naturally occurring clinical cases.

The development of effective treatments hinges on elucidating the etiology of these diseases, which likely involve a functional role for the oral microbiota and interplay of the microbiome with host genetics and environmental factors. While additional functional profiling will be necessary to pin down the role of the oral microbiota in regulating host health and disease, the work here chronicled the microbial composition of both healthy, aggressive periodontitis, and gingivostomatitis in feline oral cavities at a previously unseen taxonomic resolution. This species-level analysis of the feline oral microbiome across disease states provides a foundational roadmap for further investigation into how the mammalian oral microbiome influences debilitating inflammatory states in the oral cavity.

## Conclusion

This study reported changes in the abundance and function of the oral microbiome of cats with aggressive periodontitis (AP) and gingivostomatitis (FCGS) at high resolution and accuracy using ultra-deep shotgun metatranscriptomics. The identification of oral microbial composition in healthy, AP, and FCGS cats illustrated that species-switching underlies changes from a healthy oral cavity to inflammation of the periodontium, and concurrent inflammation of keratinized and non-keratinized epithelium. This subtle shift between healthy and disease-related microbiomes led to the proposition that function, rather than taxonomy or a singular pathobiont, contributes to disease status and could be utilized clinically. At the community level, amino acid metabolism is differentially expressed in AP and FCGS cohorts as compared to their healthy counterparts, with arginine standing out as a key amino acid regulating both host and bacterial function. Within arginine metabolism is the production of known bioactive and oral-related compounds nitric oxide and polyamines. The importance of these compounds in the context of both bacteria and host regulation suggests an interkingdom cycle wherein host pressures such as tissue deterioration seen in AP and FCGS control bacterial abundance and function in the oral cavity. This bacterial community then displays altered protein metabolism resulting in bioactive compounds that then exert effects on host inflammation and apoptotic pathways, closing the interkingdom metabolic loop and ultimately contributing to disease manifestations.

## Declarations

### Ethics approval and consent to participate

All study procedures were reviewed and approved by the University of California-Davis Institutional Animal Care and Use Committee under IACUC #22738 – “Oral Health Biobank - Repository of blood, plaque and oral tissues from cats and dogs with orodental disorders.”

### Consent for publication

## N/A

### Availability of data and material

Sequencing data generated and analyzed in this study can be found at the 100K Pathogen Project (BioProject PRJNA203445) on NCBI SRA under BioProject PRJNA1136879. Accession numbers can be found in Supplemental Table 1.

### Competing interests

The authors declare that they have no competing interests.

### Funding

This research received no external funding. MSR was supported by the National Center for Advancing Translational Sciences, National Institutes of Health, through grant number UL1 TR001860. The content is solely the responsibility of the authors and does not necessarily represent the official views of the NIH.

### Authors’ contributions

CAS conceptualized the work, analyzed and visualized the data, wrote the original draft, and edited the manuscript. MSR conceptualized the work, collected the samples, contributed to the original draft, and edited the manuscript. RP analyzed the data, visualized the data, and reviewed manuscript. CS analyzed the data and reviewed the manuscript. BCH prepared the samples for sequencing and reviewed the manuscript. AA analyzed data and reviewed the work. AB collected samples and reviewed the work. BCW conceptualized the work, contributed to the original draft, and edited the manuscript. All authors read and approved the final manuscript.

## Acknowledgements

N/A

## REFERENCES

1. Orlandi, M., et al., Impact of the treatment of periodontitis on systemic health and quality of life: A systematic review. J Clin Periodontol, 2022. 49 Suppl 24: p. 314–327.

2. Di Stefano, M., et al., Impact of Oral Microbiome in Periodontal Health and Periodontitis: A Critical Review on Prevention and Treatment. Int J Mol Sci, 2022. 23(9).

3. Pignatelli, P., et al., The Role of Fusobacterium nucleatum in Oral and Colorectal Carcinogenesis. Microorganisms, 2023. 11(9).

4. Kleinstein, S.E., K.E. Nelson, and M. Freire, Inflammatory Networks Linking Oral Microbiome with Systemic Health and Disease. J Dent Res, 2020. 99(10): p. 1131–1139.

5. Negrini, T.C., et al., Interplay Among the Oral Microbiome, Oral Cavity Conditions, the Host Immune Response, Diabetes Mellitus, and Its Associated-Risk Factors-An Overview. Front Oral Health, 2021. 2: p. 697428.

6. Soltero-Rivera, M., et al., Clinical, radiographic and histopathologic features of early-onset gingivitis and periodontitis in cats (1997-2022). J Feline Med Surg, 2023. 25(1): p. 1098612X221148577.

7. Adler, C.J., et al., Diet may influence the oral microbiome composition in cats. Microbiome, 2016. 4(1): p. 23.

8. Anderson, J.G., et al., The Oral Microbiome across Oral Sites in Cats with Chronic Gingivostomatitis, Periodontal Disease, and Tooth Resorption Compared with Healthy Cats. Animals (Basel), 2023. 13(22).

9. Older, C.E., et al., The feline cutaneous and oral microbiota are influenced by breed and environment. PLoS One, 2019. 14(7): p. e0220463.

10. Davis, E.M. and J.S. Weese, Oral Microbiome in Dogs and Cats: Dysbiosis and the Utility of Antimicrobial Therapy in the Treatment of Periodontal Disease. Vet Clin North Am Small Anim Pract, 2022. 52(1): p. 107–119.

11. Krumbeck, J.A., et al., Characterization of Oral Microbiota in Cats: Novel Insights on the Potential Role of Fungi in Feline Chronic Gingivostomatitis. Pathogens, 2021. 10(7).

12. Fried, W.A., et al., Use of unbiased metagenomic and transcriptomic analyses to investigate the association between feline calicivirus and feline chronic gingivostomatitis in domestic cats. Am J Vet Res, 2021. 82(5): p. 381–394.

13. Rodrigues, M.X., et al., The subgingival microbial community of feline periodontitis and gingivostomatitis: characterization and comparison between diseased and healthy cats. Sci Rep, 2019. 9(1): p. 12340.

14. Older, C.E., et al., Influence of the FIV Status and Chronic Gingivitis on Feline Oral Microbiota. Pathogens, 2020. 9(5).

15. Soltero-Rivera, M., S. Goldschmidt, and B. Arzi, Feline chronic gingivostomatitis current concepts in clinical management. J Feline Med Surg, 2023. 25(8): p. 1098612X231186834.

16. Rodrigues, M.X., et al., Preliminary functional analysis of the subgingival microbiota of cats with periodontitis and feline chronic gingivostomatitis. Sci Rep, 2021. 11(1): p. 6896.

17. Nakanishi, H., et al., Prevalence of microorganisms associated with feline gingivostomatitis. J Feline Med Surg, 2019. 21(2): p. 103–108.

18. Takahashi, N., Oral Microbiome Metabolism: From “Who Are They?” to “What Are They Doing?”. J Dent Res, 2015. 94(12): p. 1628–37.

19. Basic, A. and G. Dahlen, Microbial metabolites in the pathogenesis of periodontal diseases: a narrative review. Front Oral Health, 2023. 4: p. 1210200.

20. Mann, A.E., et al., Heterogeneous lineage-specific arginine deiminase expression within dental microbiome species. Microbiol Spectr, 2024. 12(4): p. e0144523.

21. Mei, F., et al., Porphyromonas gingivalis and Its Systemic Impact: Current Status. Pathogens, 2020. 9(11).

22. Sakanaka, A., et al., Fusobacterium nucleatum Metabolically Integrates Commensals and Pathogens in Oral Biofilms. mSystems, 2022. 7(4): p. e0017022.

23. Lian, J., et al., The role of polyamine metabolism in remodeling immune responses and blocking therapy within the tumor immune microenvironment. Front Immunol, 2022. 13: p. 912279.

24. Burleigh, M.C., et al., Salivary nitrite production is elevated in individuals with a higher abundance of oral nitrate-reducing bacteria. Free Radic Biol Med, 2018. 120: p. 80–88.

25. Chai, X., L. Liu, and F. Chen, Oral nitrate-reducing bacteria as potential probiotics for blood pressure homeostasis. Front Cardiovasc Med, 2024. 11: p. 1337281.

26. Coleman, J.W., Nitric oxide in immunity and inflammation. Int Immunopharmacol, 2001. 1(8): p. 1397–406.

27. Kong, N., et al., Production and analysis of high molecular weight genomic DNA for NGS pipelines using Agilent DNA extraction kit (p/n 200600). Agilent Technologies Application Note. doi, 2013. 10.

28. Basbas, C., et al., Unveiling the microbiome during post-partum uterine infection: a deep shotgun sequencing approach to characterize the dairy cow uterine microbiome. Anim Microbiome, 2023. 5(1): p. 59.

29. Garzon, A., et al., WGS of intrauterine E. coli from cows with early postpartum uterine infection reveals a non-uterine specific genotype and virulence factors. mBio, 2024. 15(6): p. e0102724.

30. Beck, K.L., et al., Monitoring the microbiome for food safety and quality using deep shotgun sequencing. NPJ Sci Food, 2021. 5(1): p. 3.

31. Bolger, A.M., M. Lohse, and B. Usadel, Trimmomatic: a flexible trimmer for Illumina sequence data. Bioinformatics, 2014. 30(15): p. 2114–20.

32. Andrews, S. Babraham Bioinformatics—FastQC A Quality Control Tool for High Throughput Sequence Data. 2010; Available from: https://www.bioinformatics.babraham.ac.uk/projects/fastqc/

33. Wood, D.E. and S.L. Salzberg, Kraken: ultrafast metagenomic sequence classification using exact alignments. Genome Biol, 2014. 15(3): p. R46.

34. Wood, D.E., J. Lu, and B. Langmead, Improved metagenomic analysis with Kraken 2. Genome Biol, 2019. 20(1): p. 257.

35. Lu, J., et al., Bracken: estimating species abundance in metagenomics data. PeerJ Computer Science, 2017. 3: p. e104.

36. Grabherr, M.G., et al., Full-length transcriptome assembly from RNA-Seq data without a reference genome. Nat Biotechnol, 2011. 29(7): p. 644–52.

37. Patro, R., et al., Salmon provides fast and bias-aware quantification of transcript expression. Nat Methods, 2017. 14(4): p. 417–419.

38. Love, M.I., W. Huber, and S. Anders, Moderated estimation of fold change and dispersion for RNA-seq data with DESeq2. Genome Biol, 2014. 15(12): p. 550.

39. Seemann, T., Prokka: rapid prokaryotic genome annotation. Bioinformatics, 2014. 30(14): p. 2068–9.

40. Huerta-Cepas, J., et al., Fast Genome-Wide Functional Annotation through Orthology Assignment by eggNOG-Mapper. Mol Biol Evol, 2017. 34(8): p. 2115–2122.

41. Hyatt, D., et al., Prodigal: prokaryotic gene recognition and translation initiation site identification. BMC Bioinformatics, 2010. 11: p. 119.

42. Eren, A.M., et al., Community-led, integrated, reproducible multi-omics with anvi’o. Nature Microbiology, 2021. 6(1): p. 3–6.

43. Cao, Y., et al., RevEcoR: an R package for the reverse ecology analysis of microbiomes. BMC Bioinformatics, 2016. 17(1): p. 294.

44. Karp, P.D., et al., Pathway Tools version 23.0 update: software for pathway/genome informatics and systems biology. Brief Bioinform, 2021. 22(1): p. 109–126.

45. Caspi, R., et al., The MetaCyc database of metabolic pathways and enzymes - a 2019 update. Nucleic Acids Res, 2020. 48(D1): p. D445–D453.

46. Soltero-Rivera, M., et al., Feline Chronic Gingivostomatitis Diagnosis and Treatment through Transcriptomic Insights. Pathogens, 2024. 13(3).

47. Das, B. and R.K. Bhadra, (p)ppGpp Metabolism and Antimicrobial Resistance in Bacterial Pathogens. Front Microbiol, 2020. 11: p. 563944.

48. Carda-Dieguez, M., R. Moazzez, and A. Mira, Functional changes in the oral microbiome after use of fluoride and arginine containing dentifrices: a metagenomic and metatranscriptomic study. Microbiome, 2022. 10(1): p. 159.

49. Chow, Y.C., et al., Implications of Porphyromonas gingivalis peptidyl arginine deiminase and gingipain R in human health and diseases. Front Cell Infect Microbiol, 2022. 12: p. 987683.

50. Battaglia, T.W., et al., A pan-cancer analysis of the microbiome in metastatic cancer. Cell, 2024.

51. Maekawa, T., et al., Porphyromonas gingivalis manipulates complement and TLR signaling to uncouple bacterial clearance from inflammation and promote dysbiosis. Cell Host Microbe, 2014. 15(6): p. 768–78.

52. Goetting-Minesky, M.P., V. Godovikova, and J.C. Fenno, Approaches to Understanding Mechanisms of Dentilisin Protease Complex Expression in Treponema denticola. Front Cell Infect Microbiol, 2021. 11: p. 668287.

53. Hyde, E.R., et al., Metagenomic analysis of nitrate-reducing bacteria in the oral cavity: implications for nitric oxide homeostasis. PLoS One, 2014. 9(3): p. e88645.

54. Lundberg, J.O., M. Carlstrom, and E. Weitzberg, Metabolic Effects of Dietary Nitrate in Health and Disease. Cell Metab, 2018. 28(1): p. 9–22.

55. Morou-Bermudez, E., et al., Pathways Linking Oral Bacteria, Nitric Oxide Metabolism, and Health. J Dent Res, 2022. 101(6): p. 623–631.

56. Rosier, B.T., et al., Nitrate reduction capacity of the oral microbiota is impaired in periodontitis: potential implications for systemic nitric oxide availability. Int J Oral Sci, 2024. 16(1): p. 1.

57. Liddle, L., et al., Variability in nitrate-reducing oral bacteria and nitric oxide metabolites in biological fluids following dietary nitrate administration: An assessment of the critical difference. Nitric Oxide, 2019. 83: p. 1–10.

58. Sturgeon, A., et al., Characterization of the oral microbiota of healthy cats using next-generation sequencing. Vet J, 2014. 201(2): p. 223–9.

59. Thomas, S., et al., Microbiome analysis of feline odontoclastic resorptive lesion (FORL) and feline oral health. J Med Microbiol, 2021. 70(4).

60. Lu, J., et al., Metagenome analysis using the Kraken software suite. Nat Protoc, 2022. 17(12): p. 2815–2839.

61. Mysak, J., et al., Porphyromonas gingivalis: major periodontopathic pathogen overview. J Immunol Res, 2014. 2014: p. 476068.

62. Simas, A.M., et al., Oral infection with a periodontal pathogen alters oral and gut microbiomes. Anaerobe, 2021. 71: p. 102399.

63. Zenobia, C. and G. Hajishengallis, Porphyromonas gingivalis virulence factors involved in subversion of leukocytes and microbial dysbiosis. Virulence, 2015. 6(3): p. 236–43.

64. Murphy, T.F. and G.I. Parameswaran, Moraxella catarrhalis, a human respiratory tract pathogen. Clin Infect Dis, 2009. 49(1): p. 124–31.

65. Verduin, C.M., et al., Moraxella catarrhalis: from emerging to established pathogen. Clin Microbiol Rev, 2002. 15(1): p. 125–44.

66. Dong, J., et al., Relationships Between Oral Microecosystem and Respiratory Diseases. Front Mol Biosci, 2021. 8: p. 718222.

67. Yu, X., et al., Moraxella occupied the largest proportion in the nasal microbiome in healthy children, which potential protect them from COVID-19. Microb Pathog, 2022. 170: p. 105685.

68. Rosier, B.T., et al., Nitrate as a potential prebiotic for the oral microbiome. Sci Rep, 2020. 10(1): p. 12895.

69. Grant, M.M. and D. Jonsson, Next Generation Sequencing Discoveries of the Nitrate-Responsive Oral Microbiome and Its Effect on Vascular Responses. J Clin Med, 2019. 8(8).

70. Wang, W., et al., The Moraxella catarrhalis nitric oxide reductase is essential for nitric oxide detoxification. J Bacteriol, 2011. 193(11): p. 2804–13.

71. Gangula, P., et al., Polybacterial Periodontal Pathogens Alter Vascular and Gut BH4/nNOS/NRF2-Phase II Enzyme Expression. PLoS One, 2015. 10(6): p. e0129885.

72. Li, Q., X. Ouyang, and J. Lin, The impact of periodontitis on vascular endothelial dysfunction. Front Cell Infect Microbiol, 2022. 12: p. 998313.

73. Zella, D., et al., Mycoplasma promotes malignant transformation in vivo, and its DnaK, a bacterial chaperone protein, has broad oncogenic properties. Proc Natl Acad Sci U S A, 2018. 115(51): p. E12005–E12014.

74. Brenner, T., A. Yamin, and R. Gallily, Mycoplasma triggering of nitric oxide production by central nervous system glial cells and its inhibition by glucocorticoids. Brain Res, 1994. 641(1): p. 51–6.

75. Drehmer, D., et al., Nitric oxide favours tumour-promoting inflammation through mitochondria-dependent and -independent actions on macrophages. Redox Biol, 2022. 54: p. 102350.

76. Baker, J.L., et al., The oral microbiome: diversity, biogeography and human health. Nat Rev Microbiol, 2024. 22(2): p. 89–104.

77. Lamont, R.J., H. Koo, and G. Hajishengallis, The oral microbiota: dynamic communities and host interactions. Nat Rev Microbiol, 2018. 16(12): p. 745–759.

78. Baratti-Mayer, D., et al., Noma: an “infectious” disease of unknown aetiology. Lancet Infect Dis, 2003. 3(7): p. 419–31.

79. Chen, C., et al., Oral microbiota of periodontal health and disease and their changes after nonsurgical periodontal therapy. ISME J, 2018. 12(5): p. 1210–1224.

80. Lyons, L.A., Cats - telomere to telomere and nose to tail. Trends Genet, 2021. 37(10): p. 865–867.

81. Fawzy El-Sayed, K.M. and C.E. Dorfer, (*) Animal Models for Periodontal Tissue Engineering: A Knowledge-Generating Process. Tissue Eng Part C Methods, 2017. 23(12): p. 900–925.

82. An, J.Y., R. Darveau, and M. Kaeberlein, Oral health in geroscience: animal models and the aging oral cavity. Geroscience, 2018. 40(1): p. 1–10.

